# SNP-based heritability estimation: measurement noise, population stratification, and stability

**DOI:** 10.1101/040055

**Authors:** Eric R. Gamazon, Danny S. Park

**Affiliations:** Division of Genetic Medicine, Department of Medicine, Vanderbilt University, Nashville, TN, USA; Academic Medical Center, University of Amsterdam, Amsterdam, The Netherlands; Department of Bioengineering and Therapeutic Sciences, University of California, San Francisco, San Francisco, CA, USA

**Author notes:** Correspondence to: Eric R. Gamazon, Ph.D. < >.

## Abstract

Siddharth Krishna Kumar^1^ and co-authors claim to have shown that “GCTA applied to current SNP data cannot produce reliable or stable estimates of heritability.” Given the numerous recent studies on the genetic architecture of complex traits that are based on this methodology, these claims have important implications for the field. Through an investigation of the stability of the likelihood function under phenotype perturbation and an analysis of its dependence on the spectral properties of the genetic relatedness matrix, our study characterizes the properties of an important approach to the analysis of GWAS data and identified crucial errors in the authors’ analyses, invalidating their main conclusions.

---

Heritability estimation using genome-wide SNP data is a fundamental research topic with profound implications for studies of the genetic architecture of complex traits. The development of a novel methodology^2,3^ in this direction has spurred studies, on a broad spectrum of complex traits, that have reinforced the view that a substantial portion of missing heritability can be accounted for by hitherto undiscovered common variants ^4,5^ and has led to substantial research that has demonstrated that certain functional categories of SNPs contribute disproportionately to the heritability of complex diseases ^6-8^. However, in a recent report ^1^, Krishna Kumar and co-authors claim to have proved that the method “may not reliably improve our understanding of the genomic basis of phenotypic variability” even when the assumptions of the method are satisfied exactly and that the heritability estimates produced are highly sensitive to the choice of sample used and to measurement errors in the phenotype. We investigated these claims by characterizing the dynamical properties of the likelihood function and identified crucial analytic errors that seriously undermine the validity of the authors’ conclusions.

## METHODS AND RESULTS

### The GREML generative model

We consider the following model (Figure 1) of the phenotype *y* (which has been simplified, as in Krishna Kumar et al., to exclude any fixed-effects):

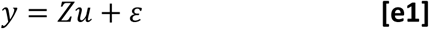

where *u* is a *Px*1 vector of random (genetic) effects, *Z* is a *NxP* (standardized genotype) matrix and *ε* is the (non-genetic) residual. Here

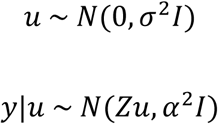

Thus, the distribution of *y* assumes the following form:

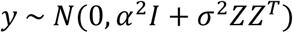

**Figure 1.**
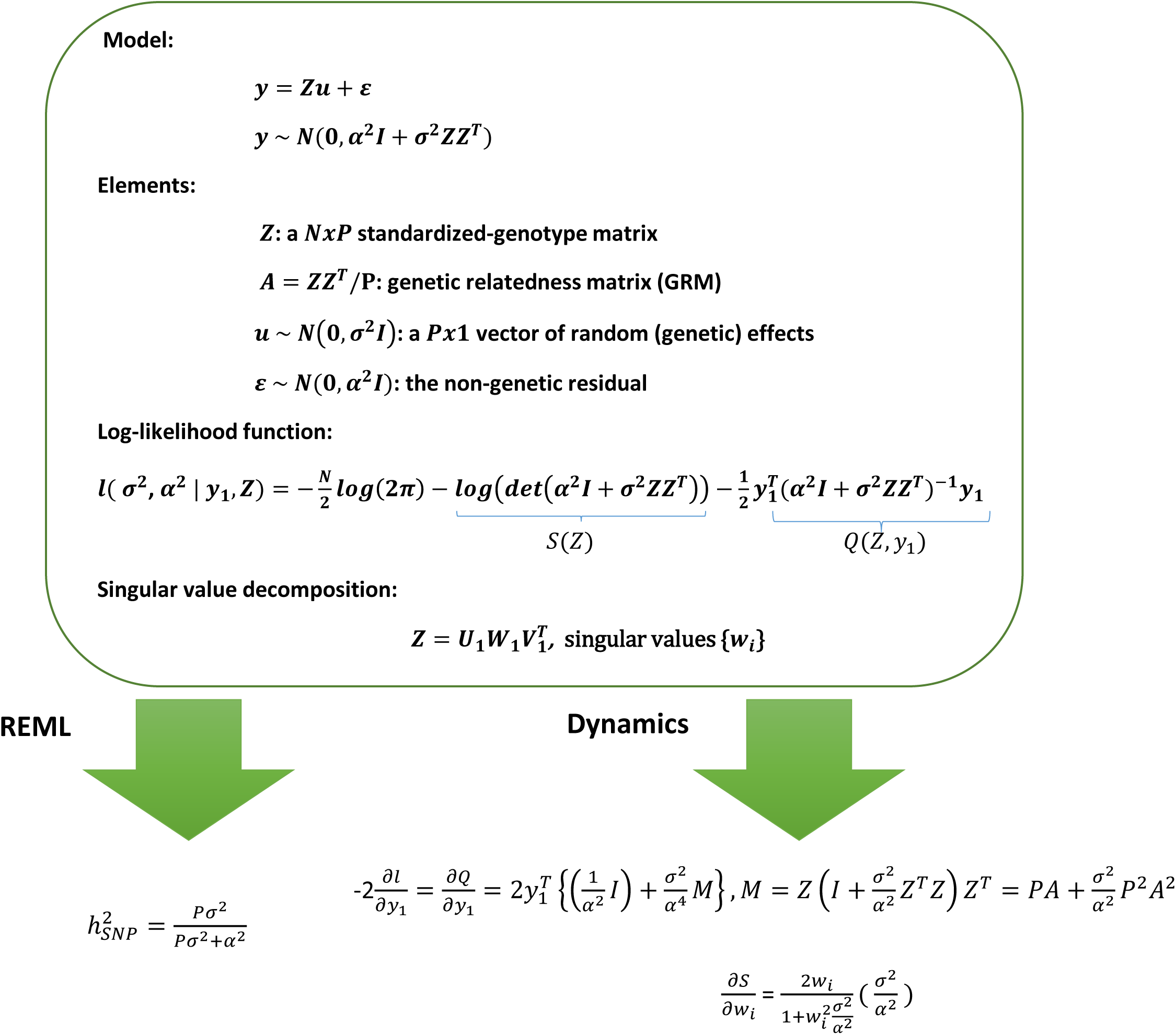
Generative model for GREML and our methodology. The GREML model (which underlies the GCTA software implementation) has been simplified here, as in the Krishna Kumar et al. study, to exclude fixed effects. Note that the phenotypic covariance decomposes into a genetic covariance and a residual covariance. The Genetic Relatedness Matrix (GRM), which quantifies the genetic similarity between pairs of individuals using the genotype data *Z*, can be written as *A* = *ZZ^T^/P*. Using Restricted Maximum Likelihood (REML), GCTA provides estimates of *σ*^2^ and *α*^2^ and thus of the SNP-based heritability: 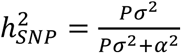. The dynamics of the log-likelihood can be investigated by considering a perturbation in the phenotype vector (e.g…, the gradient or the local curvature) or the spectral properties of the GRM.

Note that the phenotypic covariance, *var*(*y*), is the sum of a genetic covariance and a residual covariance. The Genetic Relatedness Matrix (GRM), which quantifies the genetic similarity between pairs of individuals using the genotype data *Z*, can be written as follows:

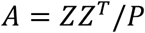

### Singularity index and induced quadratic form

We refer to the function S(Z):= log(det(*α*^2^*I* + *σ*^2^*ZZ^T^*)) as the *singularity index* (because it provides a formal test for the invertibility of the phenotypic covariance matrix *α*^2^*I* + *σ*^2^*ZZ^T^*) and refer to the function 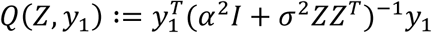 as the *induced quadratic form*. Note the log-likelihood of the observed phenotype data *y*_1_ is given by

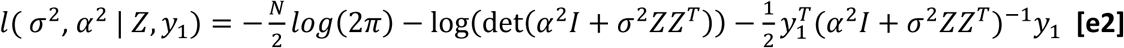

Using Restricted Maximum Likelihood (REML), GCTA estimates the variances *σ*^2^ and *α*^2^ given the observation *y*_1_, thereby providing an estimate of the SNP-based heritability:

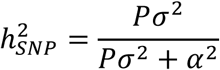

Equivalently, the log-likelihood function, now viewed as a function of *Z* and *y*_1_, can be written as a sum involving the singularity index and the induced quadratic form:

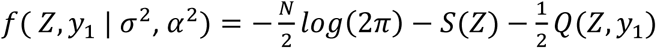

### Perturbation of the standardized genotype matrix Z and the GRM A

Because the *Z* in the GREML model is a standardized genotype matrix (wherein each entry is a function of the number of copies of the reference allele and the reference allele frequency at a SNP), this implies that there are implicit constraints on what is a *valid* perturbed genotype matrix *Z* + *perturb*(*Z*) (i.e., constraints which determine whether *Z* + *perturb*(*Z*) is a realizable or ill-defined standardized genotype matrix). A perturbation matrix *perturb*(*Z*) may generate a matrix that departs substantially from a standardized genotype matrix, yielding an ill-defined revised model. To illustrate this, if the original (e.g., independent, real and random) entries in *Z* have mean 0 and variance 1, a perturbation with elements on the primary diagonal due to the introduction of the phenotype noise *ϑ* ~ *N*(0, *τ*^2^) would preserve the mean of these elements but alter their variance, possibly quite substantially. In short, not every element of Matrices(N, P) represents a standardized genotype matrix, and not every perturbation is a reasonable one. For the same reason, a perturbation of the GRM (by an error matrix *E*, as in the authors’ equation [A17] of the Appendix) does not necessarily generate a valid (revised) GRM. (For example, the resulting perturbed GRM must be symmetric, which implies that the perturbation matrix *E* must be symmetric as well.) Furthermore, modeling the difference between the true *Z* and sample *Z* through an error matrix *F* via an additive model (*Z_sample_* = *Z_true_* + *F*) makes some very strong assumptions, including that the two matrices, *Z_true_* and *Z_sample_*, are of the same dimension (in particular, same number of variants). It is therefore more sound to evaluate the discordance between the true GRM (*GRM_true_*) and the estimated GRM (*GRM_sample_*). The impact of this discordance (arising, for example, from the imperfect tagging of causal variants ^2,9^) on the REML estimate of heritability is indeed a valid subject of research ^3^. Interestingly, this issue is related to the classic Horn’s conjecture in matrix theory (which was finally settled ^10^) on the spectrum of the sum of two Hermitian matrices and on how the eigenvalues of two Hermitian matrices constrain the eigenvalues of their sum.

### Stability analysis under phenotypic perturbation or population stratification and a framework for testing large eigenvalues

The authors evaluated the sensitivity of the likelihood function, and the resulting GREML estimate, to the GWAS data (specifically, phenotype measurement noise and population stratification). We report here crucial errors in the authors’ analyses, on which the main conclusions of the study are based. Furthermore, we highlight a methodological gap, which we address using an approach that may be of interest to future studies in population genetics and GWAS of complex traits.

We should note a random matrix theory for the Wishart product matrix *ZZ^T^* (or the GRM) generally assumes a *Z* with independent Gaussian entries, and any application in genetics must demonstrate that the relevant theoretical results apply (robustly) to a (non-Gaussian) matrix (e.g., one consisting of standardized genotype data). The authors appear to claim, clearly incorrectly and rather confusingly, for both *Z* and its symmetrization *ZZ^T^* a Wishart distribution (e.g., see pages E62 and E68 of the authors’ paper ^1^). In what follows, we will assume that *Z* is a standardized genotype matrix (and thus non-Gaussian), and Gaussian-based results that require extension to the non-Gaussian case will be explicitly stated.

#### 1. Analysis of (in)stability due to phenotype noise

The authors sought to show the instability of the induced quadratic form *Q*(*Z*, *y*_1_), and thus of the log-likelihood, by showing its sensitivity to the phenotype measurement (i.e., to a perturbation of *y*_1_). In their analysis, this conclusion follows from the instability of the spectral properties of *Z* even under a “small perturbation.” The authors used the following “equivalence” of perturbations (see equation [A10] of their Appendix A) – namely, the perturbation to the phenotype measurement and the induced perturbation of the matrix *Z*:

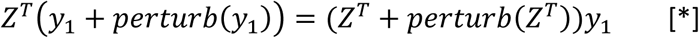

Applying the Sherman-Morrison-Woodbury identity to the third term of the log-likelihood (equation [e2]), one obtains

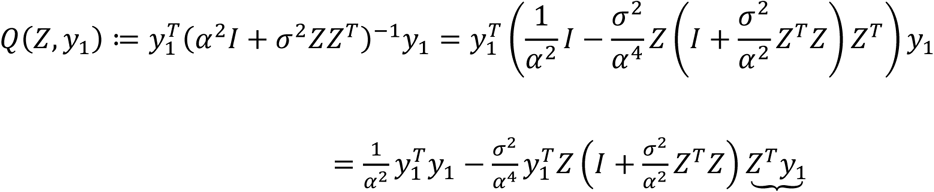

Thus, the sensitivity, assuming phenotype perturbation (equation [*]), depends not only on the factor with an underlying bracket (i.e., the spectral properties of *Z*), but also on the remaining terms (including 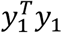). (The authors highlighted the former and, curiously, disregarded the latter.) Ignoring these remaining terms may yield invalid inferences concerning *Q*(*Z*, *y*_1_).

Importantly, *Q*(*Z*, *y*_1_) is an 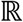-valued continuous function at every 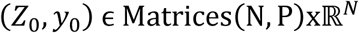, i.e.,

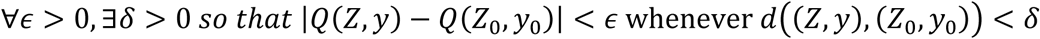

where 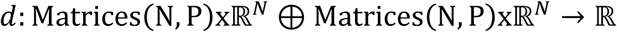 is the distance function defined by:

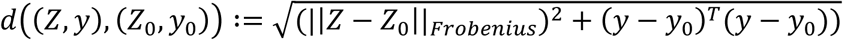

Here, for 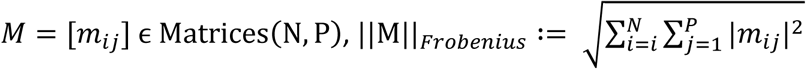. The distance function endows the set 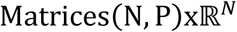 with the topology of a Euclidean space (homeomorphic to 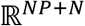) on which *Q*(*Z*, *y*), consisting of sums and products of continuous functions, is continuous. Similarly, the proper subset

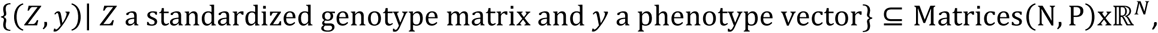

which is embeddable into 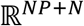 via the canonical inclusion, gets an induced subspace topology on which *Q*(*Z*, *y*) is continuous.

Given a fixed matrix *Z*, we ask how a perturbation in *y*_1_ changes *Q*(*Z*, *y*_1_). The rate of change in *Q* with respect to (the vector) *y*_1_ is given by the gradient:

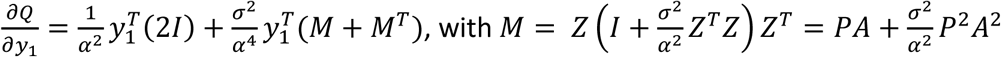

This simplifies to the following expression (by symmetry of *M*):

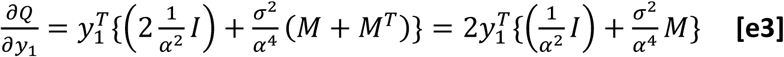

which allows us to quantify the *l*^2^-norm 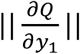 as a function of (the perturbed) *y*_1_. Because S(Z) does not depend on *y*_1_, this also gives the rate of change of the entire log-likelihood with respect to the phenotype vector (up to a constant factor). Furthermore, the expression for 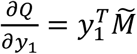 shows that *Q* ∈ *C*^1^, i.e., it is actually continuously differentiable as a function of *y*_1_. 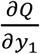 involves a second-degree polynomial in *A* and is therefore continuous as a function of the GRM. Finally, the second derivative (Hessian) matrix, which carries information about the local curvature, does not vary with the phenotype:

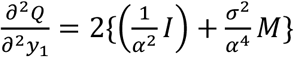

implying that higher-order derivatives do not vary with phenotype. This “curvature” matrix, along with the gradient, allows us to write a perturbation expansion, i.e., the local Taylor series expression, for the value of the log-likelihood at a perturbed value *y*_1_ + Δ*y*_1_, demonstrating the stability of the log-likelihood under phenotype measurement noise. Having ruled out phenotype perturbation as the source of the claimed instability in the likelihood function, we proceeded to characterize the dynamical properties of the likelihood under a perturbation in the genetic relatedness (see next section).

Consistent with (a) the continuity of the function *Q* in *Z* and *y*_1_, (b) the linear rate of change in *Q* with respect to *y*_1_ and (c) the well-behaved local Taylor series expansion, simulations we performed confirm the stability of the GREML estimate (Figure 2). We note that, in fact, both terms (*Q*(*Z*, *y*_1_) and S(Z)) of the log-likelihood are continuous functions at *every* 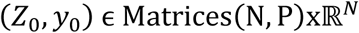.

**Figure 2.**
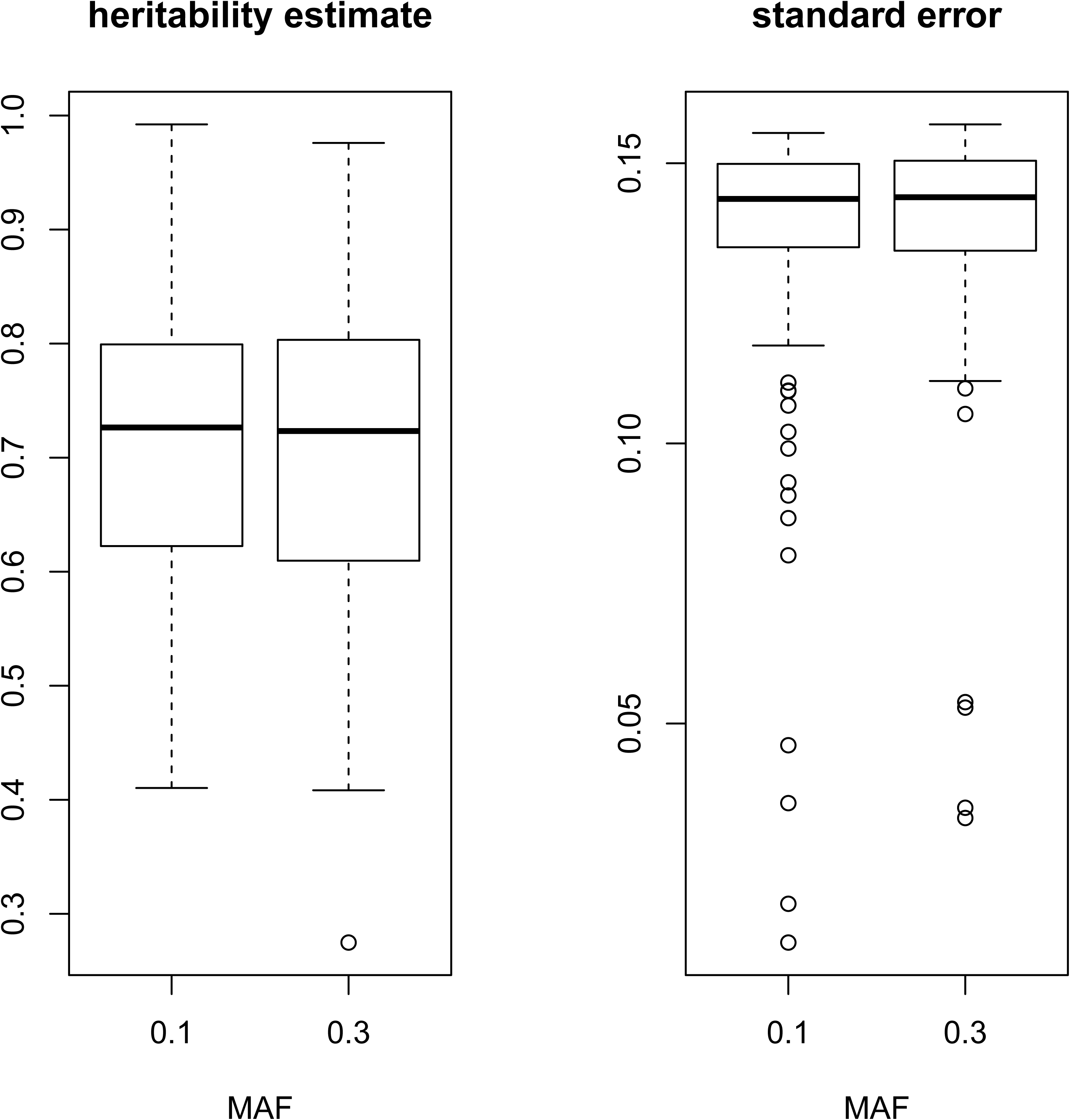
Simulation analysis to evaluate stability of the GREML estimate of heritability. We performed simulations, assuming N = 2,000 unrelated individuals, P = 50,000 independent SNPs and *h*^2^ = 0.75. For each value of the minor allele frequency (MAF) ∈ {0.10, 0.30}, we generated the matrix *Z* by drawing from the binomial distribution *Bin*(2,*maf*) and standardizing (i.e., by centering and scaling) the entries. We simulated 100 phenotypes for each MAF. The genetic effects *u* were drawn from the standard normal *N*(0,1). We used the generative model described in Figure 1 and the necessary residual to arrive at the required level of heritability. The distribution of GREML estimates for *h*^2^ and corresponding standard error is shown for each MAF.

The authors’ figure 5, which was intended to show the variation in the GREML estimates from random sampling from repeated measures of a phenotype, is *not* unexpected and, furthermore, does not empirically support the flawed theoretical argument about the instability of the log-likelihood.

#### 2. Stability of second term of log-likelihood in stratified population

Here we are interested in describing the dynamics of the likelihood function with respect to the spectral properties of the GRM in the general context (i.e., not merely when the GRM reflects population stratification). But first we consider a particular structure of genetic relatedness to evaluate the authors’ claims concerning the instability of the singularity index S(Z) under population stratification. Using the singular value decomposition (SVD) of 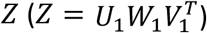 and applying the Matrix determinant lemma, one obtains the following decomposition:

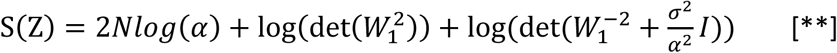

The last term of equation [**] can be written in terms of the singular values *w_i_* of *Z* as 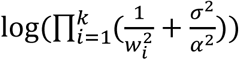. (Here we suppose that the singular values are ordered in magnitude from largest to smallest.) From this, the authors concluded (incorrectly, as we will see) that in a stratified population (for which, it is claimed, thousands of the *w_i_* are close to 0), this expression for the last term of [**] (and thus the entire expression itself) is sensitive to small changes in the values of the *w_i_*. However, one cannot show the instability of the singularity index without also considering the rest of the terms in equation [**]. Indeed, equation [**] can be rewritten as follows:

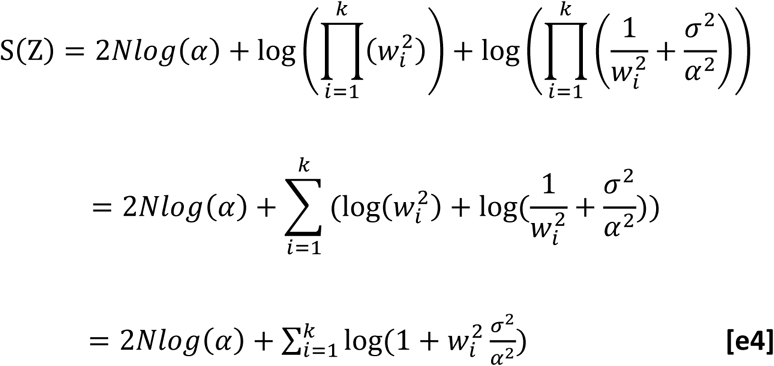

For singular values *w_i_* of *Z* that are close to 0, 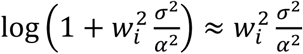(based on the Taylor series expansion). Thus, the sampling variability for near-zero singular values (from the expression for S(Z); see equation [e4]) does not arise from the terms 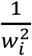 (as the authors claim), but from 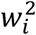. Such near-zero singular values should add little to the singularity index and closely-packed singular values (i.e., for which *w_i_* ≈ *w*, for some constant *w*) should affect S(Z) nearly similarly, and thus the claim that near-zero singular values lead to unreliable estimates of the variance explained by all SNPs (=*Pσ^2^*) remains unfounded. In contrast, very large eigenvalues (such as reflecting non-random population structure) affect the stability of the index. The rate of change of the index with respect to *w_i_*, namely 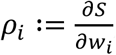, is given by the following expression:

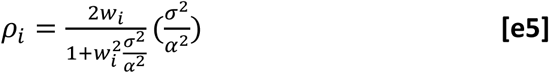

The rate-vector ρ is highly informative about the sampling behavior of the index at extreme singular values. Note that at *w_i_* = ∞, *ρ_i_* is approximately 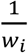. Thus, the marginal effect of increasing singular value on the index decays at infinity in a manner inversely proportional to the magnitude of the singular value. Clearly, 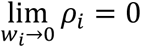, which implies that the rate of change becomes almost negligible for singular values near 0.

As we have already noted, the singularity index is also a continuous function at each 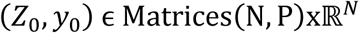 and, by projection to the first coordinate, a continuous function of the matrix *Z*. A natural question is how a perturbation (error) in the genotype matrix determines its spectral properties. The classical Weyl’s inequality^11,12^ implies that, given

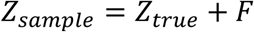

with 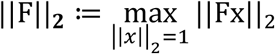, then

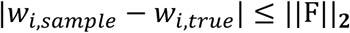

i.e., the size of the perturbation bounds the size of the resulting perturbations in singular values. A corollary of Weyl’s inequality is that the following functions:

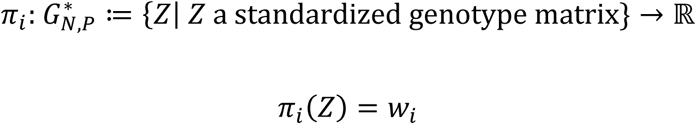

(for 1 ≤ *i* ≤ *P*) are continuous. Thus, the authors’ working premise that “a small perturbation of *Z* causes a large change in its spectral properties” (see page E67), notwithstanding the serious limitation (described above) inherent in the use of a perturbation in *Z* for this type of stability analysis, appears to conflict with this fundamental result.

If a small perturbation in *Z* implies a correspondingly small perturbation in the singular values, what can be deduced about the dynamical properties of the likelihood assuming large errors in *Z*? It is important to note that when the perturbation (i.e., ||F||_2_) is large, Weyl’s inequality, as stated (or, indeed, as in the version stated in the reference cited by the authors ^12^), sets a correspondingly large bound on |*w_i,sample_* − *w_i,true_*|. The large upper bound does not of course mean a large change in the spectral properties, but does imply *less* discrimination in our ability to distinguish between the (paired) singular values. Thus, additional machinery would be required to make accurate inferences from the singular values or the spectrum of the GRM.

#### 3. Methodological gap

What is notably missing from the authors’ analysis, given its use of the eigenvalues 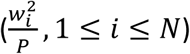 of the GRM (from the SVD) to evaluate the stability of the GREML approach, is a quantification of the degree to which the eigenvalues reflect non-random population structure versus random expectation. A large eigenvalue may well be “within null expectation,” and there is thus a need to quantify its significance. The authors failed to include a statistical framework for testing large eigenvalues. (Note this is different from the empirical distribution of the GRM eigenvalues as presented in the authors’ figure 1, which aimed to show, despite the small sample sizes considered, concordance of the data with the asymptotic behavior of eigenvalues from the Marchenko-Pastur theory.) Consideration of the null is also missing from the authors’ appropriation of the notion of an “ill-conditioned” matrix *Z*, which is defined in terms of the condition number 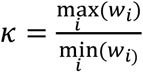, as an approach for investigating the effect on GREML estimates. In addition to these key methodological gaps, it is important to note that *κ* is a property of the matrix *Z* rather than of the GREML method. Indeed, a very large *κ* would also affect effect size estimation in simple linear regression that jointly fits multiple SNPs as fixed effects; a very large *κ* would imply that even a small change in *y* could have a destabilizing impact on the estimated SNP effect sizes and that matrix inversion would be unstable with finite-precision numbers.

The distribution of the largest eigenvalue of the Wishart matrix of a matrix *Z* with independent Gaussian entries is known ^13^. For large values of *N* and *P*, if λ denotes the largest eigenvalue, then 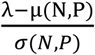 assumes the Tracy-Widom distribution ^14^; here both the centering constant μ(N, P) and the scaling constant *σ*(*N*, *P*) depend on only *N* and *P*. If the following assumptions are met for the symmetrization *ZZ^T^* = [*s_ij_*] in the GREML model (where now *Z* is the standardized genotype matrix with non-Gaussian entries):

a. the (independent real random) entries have mean 0 and variance 1
b. all moments of these random variables are finite
c. *E*(*s_ij_*)^2*m*^ ≤ *m^m^*, for some constant *m* (i.e., the distributions of the entries decay at least as fast as a Gaussian distribution)

Soshnikov’s extension theorem 15 implies that the ratio 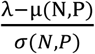, for some centering and scaling constants that depend only on *N* and *P*, converges in distribution to the Tracy-Widom distribution, just as in the Wishart case. The ratio thus provides a way to assess the significance of the largest eigenvalue of a GRM and to quantify the presence of non-random population structure in the genotype data ^16^. (For example, using the Framingham dataset presented in the authors’ figure 3, one concludes that the dataset shows extreme population stratification, p<2.2×10^-16^.) Exact expressions for the density and the moments of the distribution of the smallest eigenvalue (in terms of polynomials, exponentials and hypergeometric functions) for a matrix with independent Gaussian entries have been derived, and, interestingly, the form of this distribution depends on whether *P* − *N* is odd or even ^17^. Additionally, the work of Edelman provides a closed form for the distribution of the condition number *κ*. Indeed, for *Z* with independent standard-Gaussian entries and large *N*^17^, we can write

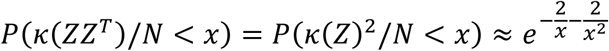

providing an asymptotic distribution for *κ*(*Z*).

The claims made by the authors concerning the stability of the GREML estimates such as through their use of the skew in singular values (such as the “Largest Singular Value” of Figure 3 and the discussion thereof in the text) are, as currently presented, statistically problematic without consideration of what is expected under the null distribution, which we characterized here.

### DISCUSSION

We investigated the dynamics of the GREML model to evaluate the dependence of the heritability estimate on phenotype perturbation and on the spectral properties of the genetic relatedness matrix. We explored the properties of the singularity index and the induced quadratic form as functions of the GRM and the phenotype. Furthermore, we derived an explicit expression for the rate of change (as well as all higher-order ones) in the log-likelihood with respect to the phenotype vector. Having ruled out phenotype perturbation as a cause of the claimed instability, we then explored the dynamical properties of the likelihood function under perturbations in the spectral properties of the GRM. In particular, we examined the sensitivity to outlier singular values, demonstrating that the authors’ claims regarding the impact of sampling variability for near-zero singular values were based on an analytic error (and assumed an incorrect view of the structure of genetic relatedness under population stratification). (It should be noted that the observation that population structure, which may be reflected in the largest eigenvalues of the GRM, may confound heritability estimation, and must thus be adjusted for, has been repeatedly discussed and investigated^18,19^.) Finally, we investigated a methodological gap in the authors’ study and highlighted an approach to address it, which may be of broad interest to methods development in population genetics and genome-wide association analysis.

## Author contributions

E.R.G. designed the study, performed the research and wrote the paper. D.S.P. performed the research. Both authors reviewed and approved the final manuscript.

## Acknowledgments

E.R.G. acknowledges support from R01 MH101820, R01 MH090937, and R01 CA157823. D.S.P. acknowledges support from K25HL121295, UCSF Krevans Fellowship, and UCSF Discovery Fellowship.

